# Relationships between chromosomal aberration breakpoints and chromosomal contacts: biophysical modeling study

**DOI:** 10.1101/089508

**Authors:** S. V. Slanina, Y. A. Eidelman, A. V. Aleshchenko, S. G. Andreev

## Abstract

Radiation-induced chromosomal exchange aberrations (CAs) are formed presumably at contacts of damaged chromosomal loci. For sparsely ionizing radiation, distribution of aberrations along the chromosome should depend on distribution of contacts, i.e. on 3D organization of the chromosome. The progress in experimental techniques for study of chromosomal contacts and for precise localization of aberration breakpoints allows to verify contacts-based mechanisms of CA formation experimentally. In the present work, the polymer model of mouse interphase chromosome 18 is developed. On this basis the experimental data on contacts and aberrations are jointly analyzed. We demonstrate high correlation between chromosomal contacts and aberrations breakpoint distributions. Possible factors and alternative mechanisms which could modify breakpoint distributions are discussed.

## Introduction

Current views on mechanisms of radiation induced chromosomal exchange aberration (CA) formation suggest misrepair of chromosomal lesions located in spatial proximity to each other [e.g. Vazquez et al. 2002, Zhang et al. 2010, Eidelman and Andreev 2015]. Since the induction of CA requires the contacts of damaged chromosomal loci, immediate (contact-first mechanism) and delayed (breakage-fist mechanism), CAs should depend of chromosomal contact pattern. Hence explanations of CA characteristics requires knowledge of the distribution of contacts within and between different chromosomes.

Until recently, any contacts-based mechanisms of CA formation could not be verified experimentally. The reasons are, first, no experimental techniques could provide adequate information about chromosomal contacts, and second, techniques capable of determining position of exchange on a chromosome (breakpoint), such as mFISH, have low resolution, (several Mbp accuracy of break position on chromosome). Everything has changed with development of Chromosome Conformation Capture technologies (3C, 4C, Hi-C, *etc*.) [Dekker et al. 2002, Lieberman-Aiden et al. 2009] and methods of accurate breakpoint determination by genome sequencing [Chiarle et al. 2011]. In [Andreev and Edel’man 1999] distributions of 4C‐ like contacts and aberration breakpoints were predicted and correlations between them were demonstrated. However, the question remained open from experimental standpoint whether this correlation/conclusion is sufficient to explain intrachromosomal aberration frequencies on the basis of information about chromosomal contacts.

The present work aims at further quantification of relationships between chromosomal contacts and intrachromosomal aberrations. The polymer model of mouse interphase chromosome 18 is developed which agrees with Hi-C data for mouse chromosome [Zhang et al. 2012]. On the basis of this model γ-induced intrachromosomal aberrations are simulated. The joint analysis of contacts and radiation-induced aberrations data [Zhang et al. 2012] demonstrates high correlation between distributions of intrachromosomal contacts and breakpoints. Possible factors and alternative mechanisms which could alter breakpoint distributions are discussed.

## Methods

The biophysical model for mouse chromosome 18 is designed on the basis of Monte Carlo technique described elsewhere [Eidelman et al. 2006, 2012]. Chromosome 18 is modeled as a heteropolymer (block-copolymer) chain of 91 megabase-sized subunits, or domains, of several types. Interphase structure of the chromosome is modeled as a result of decondensation from ultracompact mitotic chromosome. Conformational changes are determined by cis-interactions between subunits, excluded volume interactions and confining field [Eidelman et al. 2016]. Number of types, layout of all type elements and potentials for interaction between pairs of same-type and different-type subunits are determined by the iterative procedure described in [Eidelman et al. 2016].

Modeling of γ-ray induced intrachromosomal aberrations is carried out according to [Eidelman et al. 2006, 2012]. DNA double-strand breaks (DSBs) are distributed uniformly over the chromosome; they are eliminated due to repair by non-homologous end joining pathway; rates of lesion contact formation and decay are determined on the basis of calculated dynamics of all domains; if two domains with DSBs are in contact at the moment of irradiation or come in contact before the DSBs are eliminated, they interact with probability *p*, (free parameter) and the exchange aberrations is formed. Two breakpoints are assigned to positions of these domains. For comparison with the experimental data the specific characteristics used in [Zhang et al. 2012] is reproduced: distribution of breakpoints for locus 71, i.e. number of breakpoints between locus 71 and any other locus *i* as a function of *i*, after 5 Gy γ-irradiation.

## Results

### Model of mouse chromosome 18

Using Monte Carlo simulations, we develop the polymer model for mouse chromosome 18 in interphase (G1) cells. The contact map obtained for this model is shown in Fig.1 A in comparison with Hi-C data [Zhang et al. 2012] (Fig.1 B). The predicted and experimental maps are well correlated, R=0.872.

**Fig.1.**
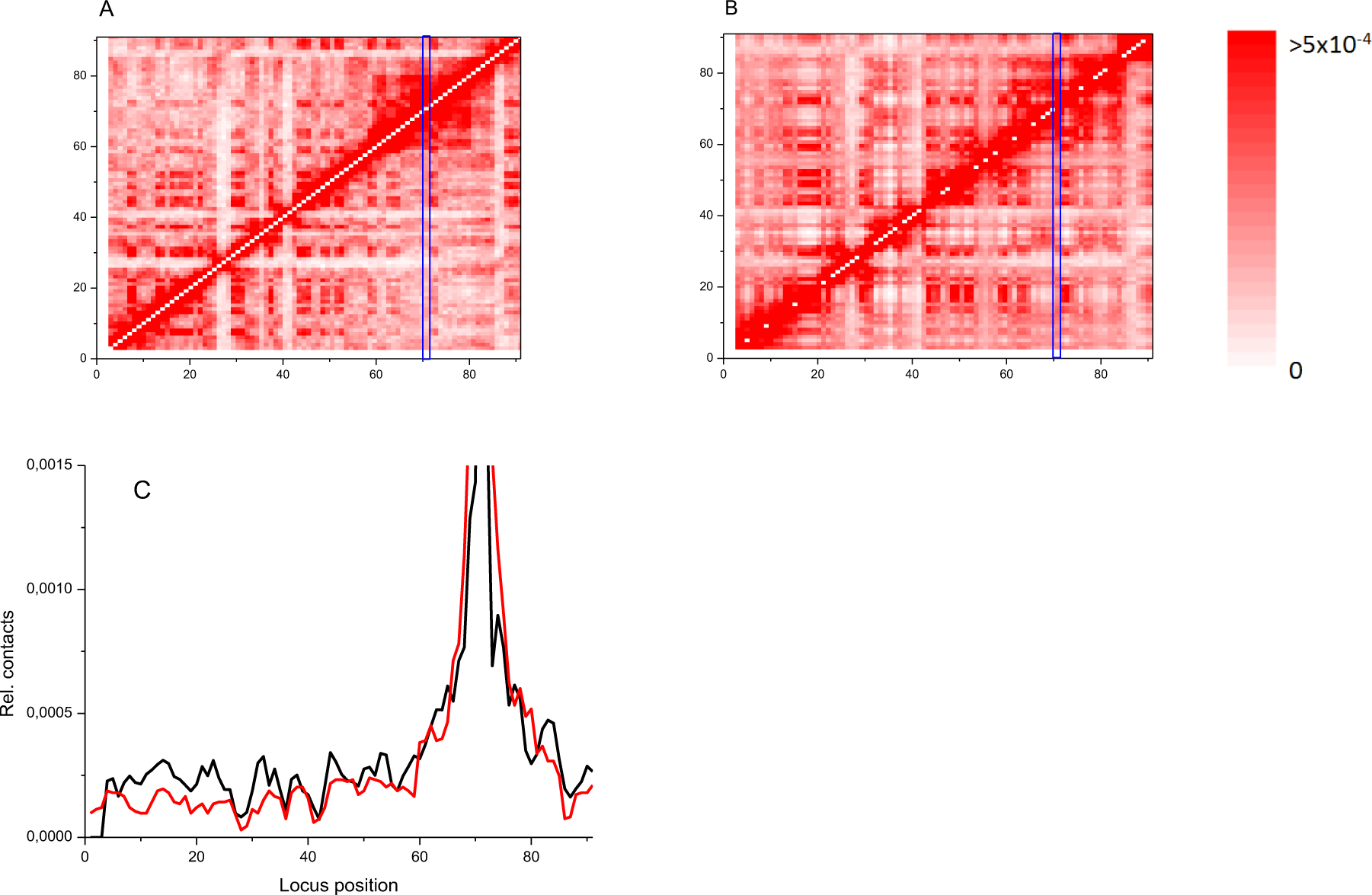
Experimental and simulated Hi-C data on mouse chromosome 18. A: contact map, simulation. B: contact map, experiment [Zhang et al. 2012]. Pearson correlation between the maps R=0.872. The blue rectangles in panels A and B show region of contacts between locus 71 and any others. C: contacts of *i*-th subunit with subunit 71. Black: experiment [Zhang et al. 2012], green: simulation. Pearson correlation coefficient R=0.916.

The contacts between locus 71 and any others (blue rectangles in Fig.1 A, B), are plotted in Fig. 1 C. These contacts are analyzed separately since it is necessary for the next stage, CA simulation. The experimental data on breakpoints available are in the format of breakpoint distribution for a locus in the region 70-71 Mbp of chromosome 18.

The simulated contact distribution, i.e. number of contacts between subunit 71 and subunit *i* as a function of *i*, agrees with the experimental distribution [Zhang et al. 2012], Pearson correlation R=0.916.

### Radiation-induced chromosomal aberrations in mouse chromosome 18

On the basis of the developed model of mouse chromosome 18 we calculated frequency of intrachromosomal aberrations following 5 Gy γ-irradiation. The simulations were made under assumption that contact-exchange probability is the same for any pair of loci. Simulated distribution of breakpoints, i.e. loci which form exchanges with locus 71, along the chromosome, is presented in Fig.2 A together with distribution of contacts of the same locus with other loci. The correlation graph between the two simulated distributions is shown in Fig.2 B.

**Fig.2.**
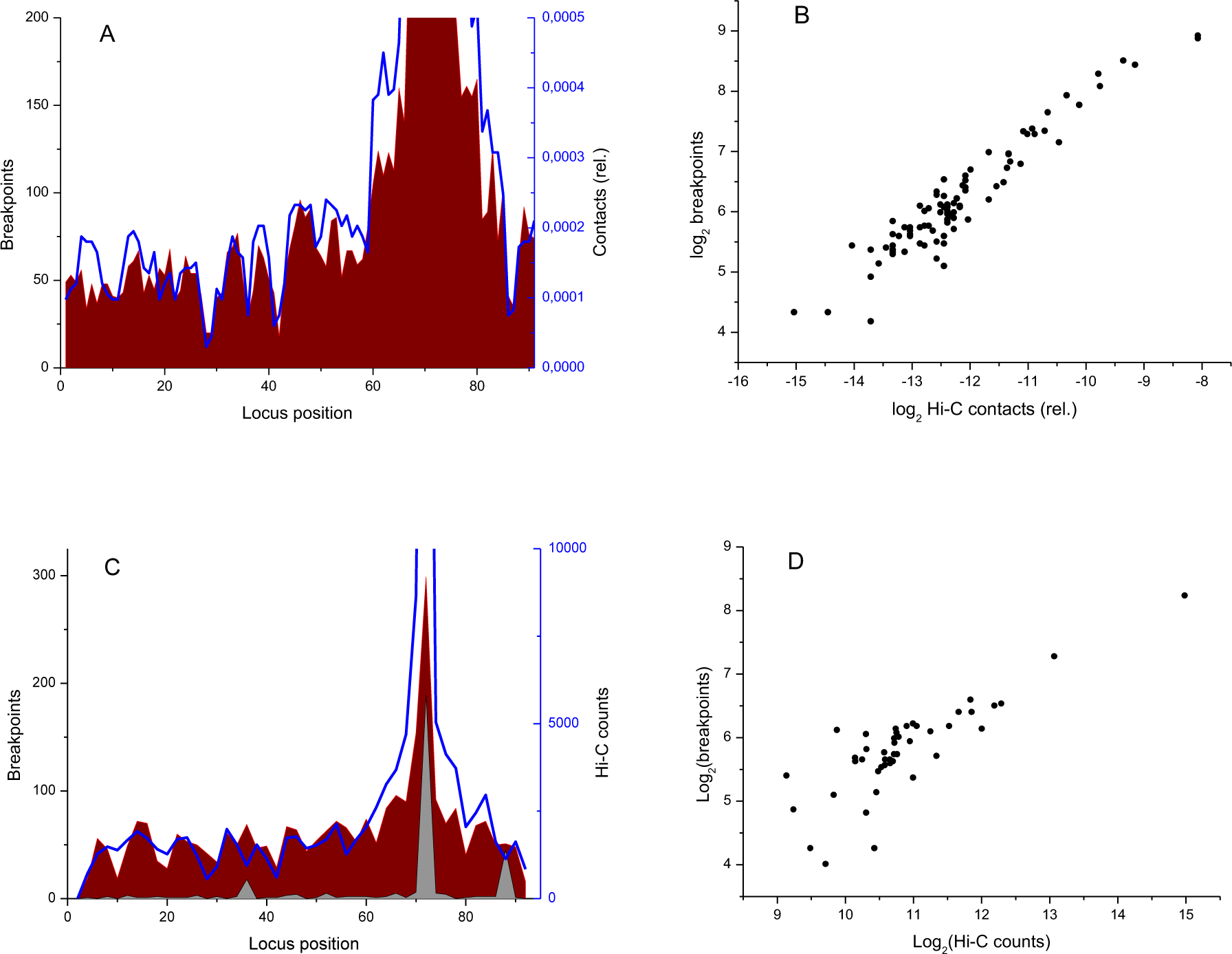
Relationship between intrachromosomal contacts and radiation induced breakpoints. Monte Carlo simulation for the polymer model of mouse chromosome 18 and experimental data. A: simulated frequency of contacts (blue line) and intrachange breakpoints (red line) between loci (*i*, 71) as a function of locus position *i*, resolution 1 Mbp. Contacts are normalized so that their sum is a fraction of contacts with locus 71 within all intrachromosmal contacts. Breakpoints are the total yield for all chromosomes considered (30 million). D=5 Gy, contact-exchange probability 0.01. B: correlation graph between contacts and breakpoints, simulation. Pearson coefficient R=0.952. C: frequency of contacts and breakpoints between loci (*i*, 71) as a function of locus position *i*, experimental data [Zhang et al. 2012], resolution 2 Mbp. Absolute counts for contacts and breakpoints are shown. Blue line: Hi-C counts; black: breakpoints for D=0, i.e. STI571-induced; red: breakpoints for D=5 Gy. D: correlation graph between contacts and breakpoints, experimental data [Zhang et al. 2012]. R=0.827.

Fig.2 C shows the experimental distributions of chromosomal contacts and breakpoints for the same locus [Zhang et al. 2012]. Correlation graph for these two distributions is shown in Fig.2 D. Distribution of exchange aberrations between subunit 71 (locus 70-71 Mbp) and other subunits of chromosome 18 are in good agreement with distribution of contacts between the same loci for experiment [Zhang et al. 2012] (Pearson correlation R=0.827) as well as for simulation (R=0.952).

The direct comparison between experimental and simulated breakpoint distributions is demonstrated in Fig.3. We conclude that the aberrations simulated on the basis of chromosome 18 computer model correlate with the experimental data [Zhang et al. 2012], R=0.770.

**Fig.3.**
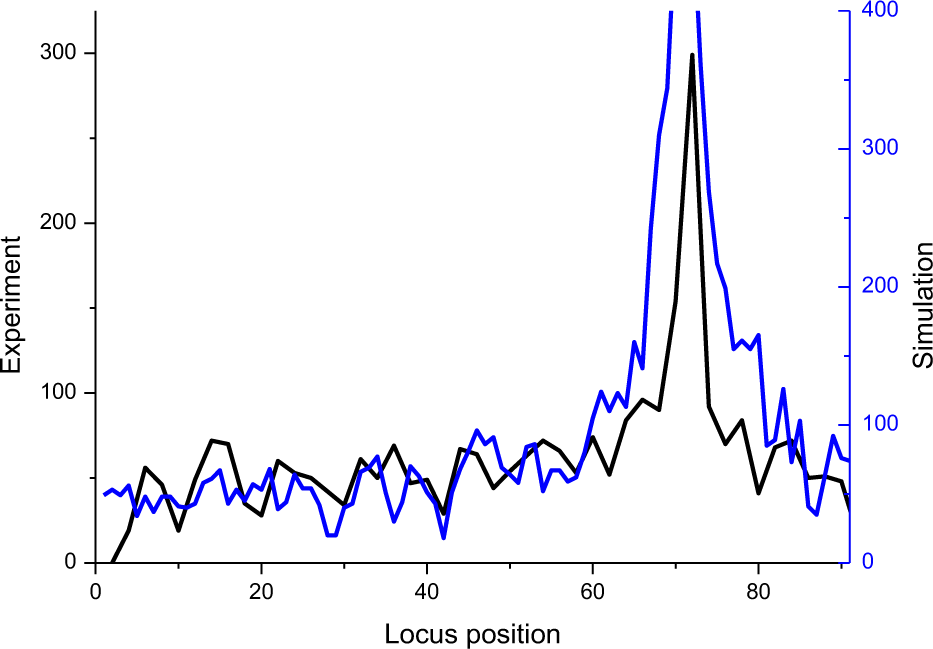
Comparison between experimental and simulated profiles of breakpoints for locus 71, D=5 Gy. Black line: experiment [Zhang et al. 2012], same data as in Fig.2 C. Blue: simulation, same data as in Fig.2 A. Pearson correlation coefficient R=0.770.

## Discussion

In the present work, the concept that chromosomal aberrations are closely related to chromosomal contacts [Wu et al. 1997, Andreev and Edel’man 1999, Savage 2000, Zhang et al. 2010] is examined on the basis of the polymer model of chromosome 18. The simple proposal is used: any contact between two damaged subunits, either being in contacts or coming in contacts, result in to exchange aberration with the constant probability *p*, contact-exchange probability.

The term “intrachromosomal” aberration used here includes all exchange aberrations formed between loci on the same chromosome. From cytogenetic point of view, this means that we analyze the whole pool of simple (centric and acentric rings, paracentric and pericentric inversions) and complex exchanges, any single exchange contributes two breakpoints, any complex exchange more than two. In [Zhang et al. 2012], aberrations were studied by genome sequencing instead of traditional cytogenetical techniques (metaphase essay, PCC). This method does not allow to distinguish aberration types but is capable to determine breakpoint positions, i.e. places on chromosomes where exchanges occurred, with high accuracy, several kbp compared to several Mbp for cytogenetic techniques. The term “translocation” used in [Zhang et al. 2012] means a breakpoint rather than a translocation as a type of interchromosomal exchange aberration in cytogenetical language.

The agreement between simulations and the experimental data [Zhang et al. 2012] both on Hi-C contacts and on CAs supports the notion made in [Andreev and Edel’man 1999] that in case of sparsely ionizing radiation which induces DNA DSBs randomly distribution of breakpoints should be determined by distribution of chromosomal contacts. In other words, the problem of aberrations comes down to the problem of chromosomal contacts.

Here we observe that the correlation between locations of chromosomal contacts and location of points where aberrations are formed in simulations (R=0.952) is higher than in experiment (R=0.827). The simple explanation is that contact-exchange probability may depend on position of specific locus on chromosome, e.g. probability may be different for loci belonging to active/inactive chromatin, early/late replicating chromatin, *etc*. Another explanation is possible contribution of the factor used in [Zhang et al. 2012] for cell synchronization, STI571 (imatinib) as a potential inducer of CAs. Breakpoint distribution for these aberrations without irradiation, i.e. what is called usually control, is highly heterogeneous and its shape does not coincide with that for radiation-induced CAs (Fig.2 C). Correction for non-zero “control” was not done in [Zhang et al. 2012], which could also contribute to relatively low correlation observed.

There is a more radical explanation of lower correlation between experimental distributions of intrachromosomal contacts and breakpoints: in addition to the simple “contact” mechanism, some other mechanisms might contribute to aberration formation. One of candidates is potential CA formation on nuclear centers [Savage 2000, Radford 2002, Eĭdel’man et al. 2014, Eidelman and Andreev 2015].

In conclusion, the polymer model of mouse chromosome 18 which agrees with Hi-C data for mouse pro-B cells is used for joint analysis of Hi-C contact data and data on distribution of radiation-induced aberration breakpoints. The analysis reveals that distribution of chromosomal contacts along the chromosome correlates with distribution of aberration breakpoints. This correlation supports the conclusion that contacts pattern of interphase chromosomes plays a significant role in γ-ray induced chromosomal aberration formation.

## Acknowledgements

This work was supported by grant 14-01-00825 from Russian Foundation for Basic Research

## References

Andreev SG, Edel’man IuA (1999) Globular model of interphase chromosome and intrachromosomal exchange aberrations. Radiats. Biol. Radioecol. 39: 10–20

Chiarle R et al. (2011) Genome-wide translocation sequencing reveals mechanisms of chromosome breaks and rearrangements in B cells. Cell 147: 107–119. doi: 10.1016/j.cell.2011.07.049

Dekker J, Rippe K, Dekker M, Kleckner N (2002) Capturing chromosome conformation. Science 295: 1306–1311. doi: 10.1126/science.1067799

Eidelman YA, Ritter S, Nasonova E, Lee R, Talyzina TA, Andreev SG (2006) Prediction of dose response for radiation induced exchange aberrations taking cell cycle delays into account. Radiat. Prot. Dosim. 122: 185–187. doi: 10.1093/rpd/ncl413

Eidelman YA, Slanina SV, Salnikov IV, Andreev SG (2012) Mechanistic modelling allows to assess pathways of DNA lesion interactions underlying chromosome aberration formation. Rus. J. Genet. 48: 1247–1256. doi: 10.1134/S1022795412120022

Eĭdel'man IuA, Slanina SV, Andreev SG (2014) Biophysical modeling of dose response for γ-ray induced complex chromosomal aberrations. Radiats. Biol. Radiobiol. 54: 140–152.

Eidelman YA, Andreev SG (2015) Computational model of dose response for low-LET-induced complex chromosomal aberrations. Radiat. Prot. Dosim. 166: 80–85. doi: 10.1093/rpd/ncv193

Eidelman Y, Slanina S, Aleshchenko A, Andreev S (2016) Chromosome interactome inferred from mitosis-G1 transition. bioRxiv doi: http://dx.doi.org/10.1101/084608

Lieberman-Aiden E et al. (2009) Comprehensive mapping of long range interactions reveals folding principles of the human genome. Science 326: 289–293. doi: 10.1126/science.1181369

Radford IR (2002) Transcription-based model for the induction of interchromosomal exchange events by ionizing irradiation in mammalian cell lines that undergo necrosis. Int. J. Radiat. Biol. 78: 1081–1093. doi: 10.1080/0955300021000034684

Savage JRK (2000) Proximity matters. Science 290: 62–63. doi: 10.1126/science.290.5489.62

Vazquez M, Greulich-Bode KM, Arsuaga J, Cornforth MN, Brückner M, Sachs RK, Hlatky L, Molls M, Hahnfeldt P (2002) Computer analysis of mFISH chromosome aberration data uncovers an excess of very complicated metaphases. Int. J. Radiat. Biol. 78: 1103–1115. doi: 10.1080/09553000210166354

Wu H, Durante M, Sachs RK, Yang TC (1997) Centric rings, acentric rings and excess acentric fragments based on a random-walk interphase chromosome model. Int. J. Radiat. Biol. 71: 487–496. doi: 10.1080/095530097143815

Zhang Y, Gostissa M, Hildebrand DG, Becker MS, Boboila C, Chiarle R, Lewis S, Alt FW (2010) The role of mechanistic factors in promoting chromosomal translocations found in lymphoid and other cancers. Adv. Immunol. 106: 93–133. doi: 10.1016/S0065-2776(10)06004-9

Zhang Y, McCord RP, Ho YJ, Lajoie BR, Hildebrand DG, Simon AC, Becker MS, Alt FW, Dekker J (2012) Spatial organization of the mouse genome and its role in recurrent chromosomal translocations. Cell 148: 908–921. doi: 10.1016/j.cell.2012.02.002

